# sdAbs-LLM: Generative Large Language Models For *de novo* Antibody Design and Agentic Evaluation

**DOI:** 10.64898/2026.04.18.716776

**Authors:** Delower Hossain, Fuad Al Abir, Sixue Zhang, Jake Y. Chen

## Abstract

Despite major advances in computational antibody engineering, no systematic comparison of modern open-source LLM backbone families for antibody sequence generation exists, nor is it known whether architectural differences matter at compact model scales. In this study, five compact transformer variants inspired by prominent open-source LLM families (Llama-4, Gemma-3, DeepSeek-V3, Mistral 7B, and NVIDIA Nemotron-3) were customized and trained from scratch for de novo VH single-domain antibody (sdAb) design. All five models were pretrained from scratch on 15 million sequences from the Observed Antibody Space (OAS) database. Pretraining yielded uniformly high generative fidelity across architectures: sequence diversity 0.507–0.516 (CV=0.8%), uniqueness approaching 1.0, and novelty 0.925–0.977 (CV=2.2%). The models were subsequently fine-tuned on disease-stratified repertoires spanning SARS-CoV-2 (n=4,688), HIV (n=430), HER2 (n=22,778), and Ebola virus (n=2,868). Structural assessment of top-ranked candidates of those case studies via AlphaFold-2, Boltz-2, RoseTTAFold-2, and ESMFold produced mean pLDDT scores of 92.88±1.54 to 93.77±2.16, with no statistically significant inter-model differences (Kruskal–Wallis H=2.06, p>0.05; N=100), indicating no statistically detectable difference was observed across architectures at this compressed scale in a single-seed experiment, suggesting that generative capacity at this parameter regime is primarily determined by training data and model scale rather than family-specific design elements at this scale. Computational docking yielded predicted binding free energies of −36.34 to −65.60 kcal/mol; independent biological rigor validation through IMGT-defined CDR-H3 extraction, BLASTp novelty assessment, and NetMHCIIpan 4.3 MHC-II immunogenicity profiling collectively confirmed antigen-binding loop novelty (CDR-H3 identity 0–29% to closest database hits), germline-consistent humanness (77–90% VH germline content), and immunogenically silent antigen-binding surfaces with no strong MHC-II binders detected across CDR regions in any candidate. We further introduce a proof-of-concept agentic evaluation pipeline leveraging the Model Context Protocol (MCP) with Claude Sonnet 4.6, enabling automated structural profiling and candidate prioritization across disease targets.

## 1 Introduction

Antibodies are immunoglobulin proteins secreted by B cells that selectively recognize foreign molecules, known as antigens, to initiate a targeted immune response. Each antibody is composed of paired heavy and light chains, whose conserved framework regions flank three hypervariable Complementarity Determining Region (CDR) loops that directly mediate antigen binding. The human immune system generates an estimated trillion of distinct antibody specificities even before exposure to an antigen [1]. Humanized monoclonal antibodies have achieved widespread therapeutic success, demonstrating efficacy across diverse conditions, including oncology, autoimmune diseases, and infectious disorders, and now represent the leading class of approved drug modalities.

More than 160 therapeutic antibodies have been approved worldwide since the first monoclonal antibody drug, muromonab-CD3 (OKT3), was permitted by the US Food and Drug Administration in 1986, and the global market is projected to exceed US$445 billion within the next five years [2]. Emerging modalities, including bispecific antibodies and antibody drug conjugates (ADCs), continue to expand the therapeutic space and sustain demand for accelerated discovery pipelines [3]. Within the broader antibody landscape, VH single-domain antibodies (sdAbs, also termed nanobodies when derived from camelid species) have emerged as a therapeutically relevant modality with distinct advantages, including small size (~12–15 kDa), enhanced tissue penetration, and amenability to expression in recombinant systems. The FDA approval of caplacizumab in 2019 validated the therapeutic potential of sdAbs, and numerous candidates are in clinical development [6]. This study targets de novo VH sdAb design, not full-length IgG. That choice constrains the design space to a single VH domain that must function as an autonomous binding unit.

AI-driven antibody discovery is reshaping modern computational biotherapeutics [5][6][7][8][9][10][11]. A landmark advance in this domain is AlphaFold, a deep learning system based on a multiple sequence alignment attention architecture (Evoformer), developed by John Jumper at DeepMind that predicts highly accurate three-dimensional protein structures directly from amino acid sequences [4]. The NVIDIA Nemotron-3 and DeepSeek-V3 remain largely unexplored in this domain.

To bridge this gap, the present study adopts compact backbone configurations inspired by these state-of-the-art models and trains them from scratch for de novo VH sdAb design. Due to computational constraints and domain realism, the models evaluated here are substantially reduced in scale compared to their original published configurations (Table 2). Models are pre-trained on 15 million benchmark antibody sequences from the OAS dataset and subsequently fine-tuned across four immunologically relevant disease targets: Ebola virus, Human Immunodeficiency Virus (HIV), Human Epidermal Growth Factor Receptor 2 (HER2), and Severe Acute Respiratory Syndrome Coronavirus 2 (SARS-CoV-2). The key contributions of this article are summarized as follows:

- Introduced compact LLM Variants: Five compact LLM-inspired models trained from scratch generate VH sdAb candidates (~8h/model on A100 80GB).
- Rigor Multi-Disease Validation: Generated antibodies evaluated across multiple diseases with statistical analysis.
- Agentic Evaluation: MCP + Claude pipeline orchestrates AlphaFold-2, Boltz-2, RoseTTAFold-2, and ESMFold for automated structural evaluation.

## 2. Related Works

Transformer-based generative Large Language Models (LLMs) for antibody design have been explored, with a focus on BERT, GPT, ESM, and LLaMA architectures. In 2022, Olsen et al. [14]and Akbar et al. [15]introduced AbLang and Ab-BERT, respectively, establishing foundational BERT-style masked language models for antibody sequence modeling. Subsequently, GPT-based autoregressive models were developed to enhance generative capabilities, including AB-Gen [18] and IgLM [19], while Zhang et al. [16] and Liu et al. [20] presented AB-GPT and p-IgGen, respectively, further growing commercial momentum further underscores the field’s promise: in December 2023, AstraZeneca and AbbVie each signed partnerships exceeding $200 million with Absci and BigHat Biosciences, respectively, to accelerate next-generation therapeutic development through AI-driven antibody design platforms [11]. Despite significant progress, key challenges remain. Existing AI approaches, including GAN-LLM hybrid frameworks [11], graph-based models such as AbFlex [12]and MEAN [13], and generative language (GenAI) models, including ABLang [14], AB-BERT [15], and Ab-GPT [16], exhibit limitations in generalizability and computational resource utilization. In contrast, the potential of modern transformer-based architectures, including LLaMA-4, Gemma-3, Mistral 7B, demonstrating the potential of GPT-style frameworks for generating full-length antibody sequences. In parallel, ESM-based models have been employed to generate antibodies. For example, PALM-H3 [22]targets CDR3 region generation in SARS-CoV-2 antibodies, leveraging the ESM-2 backbone in conjunction with RoFormer attention mechanisms, and S2LM [21]similarly utilizes ESM-2 embeddings to inform structure-aware sequence modeling. More recently, in 2025, NanoAbLLaMA [23]applied fine-tuning on LLaMA-2 to enable nanobody sequence generation, although LLaMA has since introduced more updated architectures. However, despite their promise, these approaches have notable limitations. Fine-tuned models like NanoAbLLaMA may retain irrelevant parameters and yield moderate pLDDT scores, while models such as S2LM may be slow for high-throughput screening. Across methods, comprehensive silico evaluation—including molecular dynamics studies and benchmarking metrics such as novelty, diversity, uniqueness, perplexity, and Jaccard similarity—is often missing, limiting rigorous assessment of generative performance. Many prior approaches also lack comprehensive case studies demonstrating practical applicability for disease-driven antibody library design.

### Contextual comparison with prior antibody generative models

This study does not perform direct experimental comparisons with prior antibody generative models but provides a literature-based contextual comparison (Table 1). Models such as IgLM [19], NanoAbLLaMA [23], and PALM-H3 [22] report structural confidence or perplexity metrics for generated antibodies. The compact models in this work achieve a mean pLDDT of 93.14 ± 2.18 for the top 2% candidates, indicating strong structural confidence. However, differences in antibody formats and evaluation protocols limit direct comparison. Structure-first diffusion methods like Rfdiffusion [2] and DiffAb follow a different design paradigm and are not directly comparable.

**Table 1:** Literature comparison of antibody generative models (values extracted from published papers)

| Method | Architecture | Format | Training Data | Reported pLDDT | Diversity | Novelty | Key Limitation |
| --- | --- | --- | --- | --- | --- | --- | --- |
| IgLM [19] | GPT-2 | Full IgG | OAS (~2M) | Not reported | Moderate | High | No structural eval |
| NanoAbLLaMA [23] | LLaMA-2 (FT) | Nanobody | ~15K nanobodies | 70–88 (top) | Moderate | High | Moderate pLDDT |
| PALM-H3 [22] | ESM-2 + RoFormer | CDR-H3 only | SARS-CoV-2 abs | 75–90 (top) | High | High | CDR-only, one target |
| AB-Gen [18] | GPT-2 | Full VH/VL | OAS subset | Not reported | Moderate | High | No 3D evaluation |
| LLM-Ab (Ours) | Compact LLM variants | VH sdAb | OAS (15M) + disease | 93.14 ± 2.18 (top 2%)<br>Very High | 0.51 | Very High | Top-candidate only; no baselines. |

Note: Published values are approximate and extracted from respective papers under potentially different evaluation conditions. Direct numerical comparison requires identical evaluation protocols and is provided here only for rough contextual reference. The pLDDT values for this study reflect the top 20 of 1,000 candidates and are not directly comparable to studies reporting full-distribution statistics.

**Table 2:** Architectural comparison of candidate (LLMs) for antibody design and development.

| Features | Llama (4) | Gemma -3 | Mistral | Nemotron-3 | DeepSeek -V3 |
| --- | --- | --- | --- | --- | --- |
| **Architecture** | MoE | Decoder-only | Decoder-only | Decoder-only | MoE |
| **Positional Encoding** | iRoPE | Rotary (RoPE) | Rotary (RoPE) | Rotary (RoPE) | Rotary (RoPE) |
| **Normalization** | RMS Norm | RMS Norm | RMS Norm | RMS Norm | RMS Norm |
| **Attention Mechanism** | Grouped Query (GQA) | Grouped Query (GQA) | Sliding Window (SWA) | Grouped Query (GQA) | Multi-head Latent (MLA) |
| **Memory Efficiency** | KV Cache | KV Cache | Rolling Buffer Cache | KV Cache | MLA |
| **Activation** | SwiGLU | GeGLU | SwiGLU | SwiGLU | SwiGLU |

MoE refers to Mixture of Experts as specialized expert layers per token to improve scalability and efficiency, while iRoPE stands for interleaved Rotary Positional Encoding.

## 3. Methodology

### 3.1 Dataset

The Observed Antibody Space (OAS) [24]dataset was used to pre-train on 15 million antibody sequences to capture core sequence patterns and biological semantics. The dataset provides full amino acid sequences, chain types, complementarity-determining regions (CDRs), and framework regions (FWRs), and species class. The pretrained models were then fine-tuned on from disease-associated repertoires antibody datasets, including SARS-CoV-2 (n = 4,688) [25], Human Immunodeficiency Virus (HIV; n = 430) [26], Human Epidermal Growth Factor Receptor 2 (HER2; n = 22,778) [27], and Ebola virus (n = 2,868) [28].

### 3.2 Data pipeline

We implemented a fixed-vocabulary tokenizer for antibody sequences using the Hugging Face Tokenizers library with a BPE backend. The vocabulary comprises 20 canonical amino acid residues and three special tokens ([PAD], [BOS], [EOS]). Unlike natural language tokenization, no subword merging was performed; each amino acid constitutes a biological atomic unit, making single-residue tokenization biologically appropriate. Sequences were tokenized with [BOS] and [EOS] boundary markers, then truncated or padded to a maximum length of 512 tokens and converted to PyTorch tensors. Fine-tuning adjusts the generative distribution toward disease-associated repertoire statistics but does not condition on antigen structure; generated candidates statistically resemble disease-associated antibodies but binding to the intended antigen is not guaranteed. Note on test set design. The current training used a 90/10 training/validation split without a held-out test set. Reported perplexity values are validation metrics; generalization to unseen sequences is not formally evaluated. The extended phase should adopt an 80/10/10 train/validation/test split.

### 3.3 Architecture and Hyperparameters Setup (Compact Transformer Variant Suite: Llama-4, Gemma-3, Mistral, DeepSeek-V3, Nemotron-3)

Large language models have become central to modern AI, especially in multi-agent systems that rely on LLMs for orchestration, novel content generation, and reasoning attributes. We adopted compact configurations inspired by leading open-source LLM backbones: Llama-4 [29], Gemma-3 [30], DeepSeek-V3 [31], Nemotron-3 [32], and Mistral 7B [33]. Table 2 summarizes the key architectural features and mechanisms of the original published systems. Despite variations among these models, several common design elements are observed. Most architectures follow a Transformer-based decoder-only framework, while Llama-4 and DeepSeek-V3 incorporate Mixture-of-Experts (MoE) layers to improve scalability and parameter efficiency. For positional representation, most models use rotary-based positional encoding (RoPE), whereas Llama-4 uses an improved variant, iRoPE. All models apply RMSNorm for normalization due to its computational efficiency and training stability.

**Figure 1:**
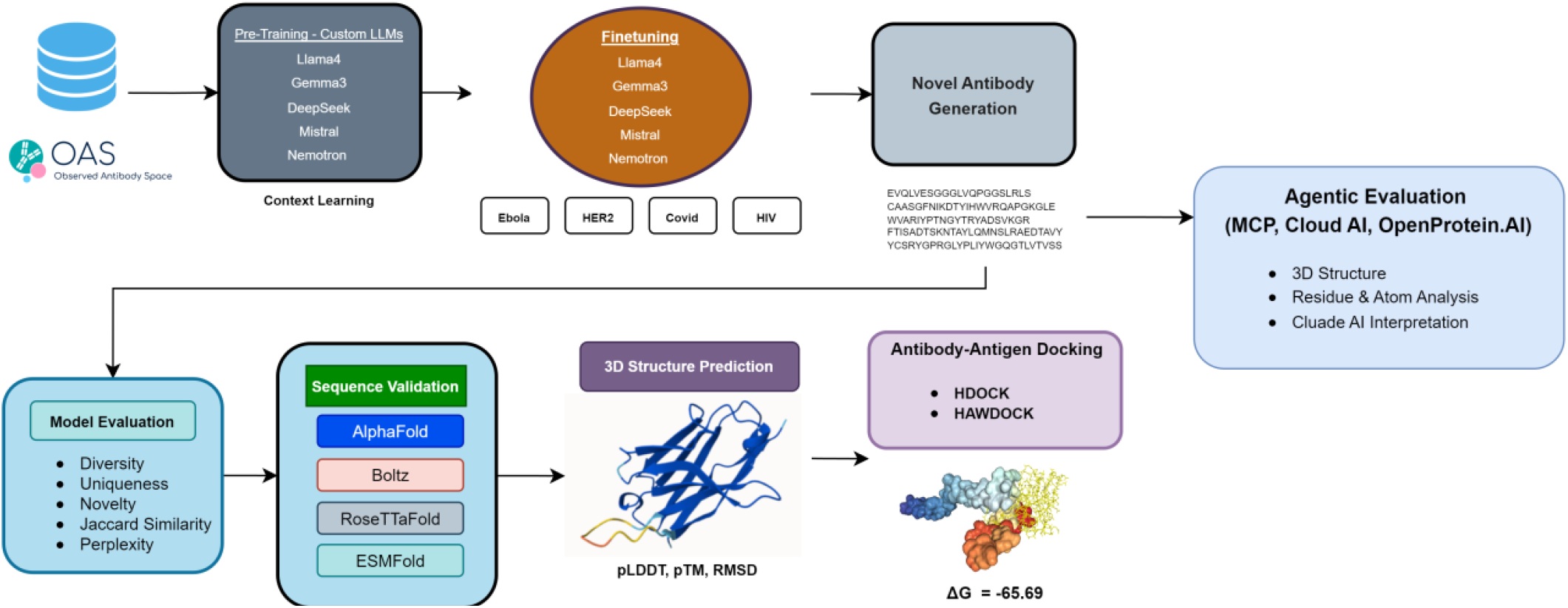
End-to-end computational pipeline for de novo antibody design and agentic evaluation. The framework integrates large language model fine-tuning on the Observed Antibody (OAS) database, generative sequence sampling across four pathogen targets, multi-tool 3D structure prediction, and automated agentic evaluation via MCP-enabled Claude AI services.

#### Critical note on model scale

Although the selected LLM families vary substantially in original scale, from Gemma-3 (1B) to Llama-4 (109B), all architectures were reduced to a uniform compact configuration (6 layers, 384 hidden dimensions, 4 attention heads, ~2–10M parameters), retaining only family-specific design elements: activation functions, normalization settings, and attention configurations. At this scale, mechanisms such as MoE routing, Sliding Window Attention, and Multi-head Latent Attention cannot fully express their intended computational advantages. Results should therefore be interpreted as an assessment of whether family-specific inductive biases yield measurable differences in antibody generation quality under a compressed regime, rather than a comparison of the full-scale published architectures. Importantly, despite extreme parameter reduction, biological validation confirmed these compact architectures successfully learned antibody biological grammar, evidenced by germline-consistent frameworks (77– 90% VH germline content), novel antigen-binding loops (CDR-H3 identity 0–29%), and immunogenically silent CDR regions.

#### The hyperparameter configurations (Appendix A)

All five architectures shared identical hyperparameters (AdamW, lr=2×10^−5^, batch size=128, gradient accumulation=4 steps, cross-entropy loss, seed=42) to isolate architectural effects. Models accept amino acid sequences up to 512 tokens with boundary tokens, and training was logged via Weights & Biases. Each variant was pretrained on OAS-15M for three epochs (~8 hours, single NVIDIA A100 80GB GPU), with architectural differences limited to activation functions (ReLU^2^ for Nemotron, SiLU for Llama/Mistral/DeepSeek, GELU (tanh) for Gemma) and normalisation parameters. Fine-tuning used identical settings with convergence-adjusted epochs: SARS-CoV-2 (80), HIV (60), HER2 (25), and Ebola (5).

#### Epoch selection rationale

Fine-tuning epochs (60-250) were selected by weights and bias (WB) online monitoring of validation loss trajectories (Appendix B), rather than by automated early stopping. The non-monotonic relationship between dataset size and epoch count (COVID-19: 80 epochs on n = 4,688; HER2: 25 epochs on n = 22,778) reflects differences in convergence speed driven by intrinsic sequence diversity rather than dataset size alone. Automated early stopping with patience-based criteria should be adopted in future work. Automated early stopping with patience-based criteria should be adopted in the future.

#### Novel generation measurement scope

Novelty was defined as the absence of exact matches between generated sequences and the respective training set, a conservative measure that does not account for near-duplicates or high-identity subsequences; BLAST-based assessment (>90% identity threshold) is reported in the results section. For each disease target, n=1,000 VH sdAb candidates were generated and evaluated across five sequence-level metrics: diversity, uniqueness, novelty, Jaccard similarity, and perplexity (full results in Appendix H).

#### Docking methodology and limitations

Antibody structures predicted by AlphaFold-2 were saved as PDB files. The matching antigen structures were procured from the Protein Data Bank and cleaned by removing water, ligands, and duplicate chains. For instance, in an HIV study, the gp120 envelope glycoprotein was used as an antigen. If known, the antigen’s binding region was identified to help guide docking. The cleaned antigen acted as the receptor, while the AlphaFold-predicted antibody served as the ligand, and both were submitted to HDOCK for rigid-body docking to evaluate antigen–antibody interactions. HDOCK and HawDOCK use rigid-body docking and ignore antibody flexibility, so the reported ΔG values are only rough computational estimates. No flexible refinement or negative controls were used, and comparisons with experimental crystal structures are only in silico; future work should dock reference antibodies through the same pipeline for fair comparison. 3.5 Agentic Evaluation:

### 3.4 Agentic Evaluation (The Proof-of-Concept)

Additionally, study presented a proof-of-concept integration of the Model Context Protocol (MCP) as an orchestration layer that connects large language models (e.g., Claude Sonnet 4.6) with structural biology APIs (OpenProtein.AI) and bioinformatics tools to enable automated, agentic antibody evaluation. Implemented via FastMCP, the framework dynamically coordinates multiple structure prediction engines (AlphaFold2, ESMFold, Boltz-2, RoseTTAFold) alongside libraries such as Biopython, modlamp, pfeature, and mhcflurry to perform end-to-end analysis, including 3D structure prediction, residue-level characterization, binding interaction modeling, and extraction of confidence metrics (e.g., pLDDT, PAE). This unified system supports multi-scale interpretation from sequence to structure to function; however, it remains an engineering demonstration and has not yet been formally benchmarked against expert-driven or scripted pipelines in terms of accuracy, efficiency, or robustness, making systematic validation an important direction for future work.

## 4. Result

### 4.1 de novo antibody design and 3D Structure Evaluation (Pre-Trained LLMs from Scratch)

This study leverages the five variants of compact LLM architectures described in Section 3.3. Table 3 summarizes the performance evaluation report, in which Llama4-Ab and Mistral3-Ab achieved the highest uniqueness (0.999) and novelty (0.977), whereas Nemotron3-Ab yielded the highest diversity (0.513) (Table).

**Table 3:**
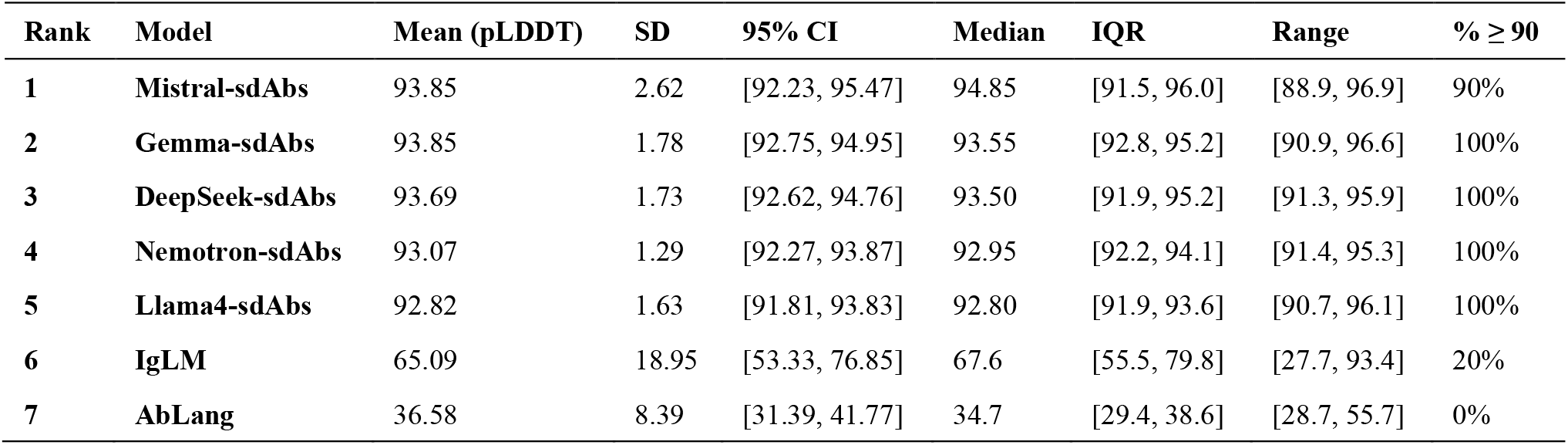
Ranked pLDDT Statistical Analysis.

| Rank | Model | Mean (pLDDT) | SD | 95% CI | Median | IQR | Range | % $\geq 90$ |
| --- | --- | --- | --- | --- | --- | --- | --- | --- |
| 1 | Mistral-sdAbs | 93.85 | 2.62 | [92.23, 95.47] | 94.85 | [91.5, 96.0] | [88.9, 96.9] | 90% |
| 2 | Gemma-sdAbs | 93.85 | 1.78 | [92.75, 94.95] | 93.55 | [92.8, 95.2] | [90.9, 96.6] | 100% |
| 3 | DeepSeek-sdAbs | 93.69 | 1.73 | [92.62, 94.76] | 93.50 | [91.9, 95.2] | [91.3, 95.9] | 100% |
| 4 | Nemotron-sdAbs | 93.07 | 1.29 | [92.27, 93.87] | 92.95 | [92.2, 94.1] | [91.4, 95.3] | 100% |
| 5 | Llama4-sdAbs | 92.82 | 1.63 | [91.81, 93.83] | 92.80 | [91.9, 93.6] | [90.7, 96.1] | 100% |
| 6 | IgLM | 65.09 | 18.95 | [53.33, 76.85] | 67.6 | [55.5, 79.8] | [27.7, 93.4] | 20% |
| 7 | AbLang | 36.58 | 8.39 | [31.39, 41.77] | 34.7 | [29.4, 38.6] | [28.7, 55.7] | 0% |

The three-dimensional (3D) structure of antibodies is crucial because it determines the spatial arrangement of complementarity-determining regions (CDRs) that govern antigen recognition and binding specificity. The top 20 candidates (2% of 1,000) were selected for AlphaFold-2 structure prediction, representing the high-quality tail of the generated sequences. The best structure per disease was further validated using AlphaFold [34][35], Boltz-2 [36], RoseTTAFold-2 [37] [38], and ESMFold [39], and rigid-body docking with HDOCK [40], and HawDOCK [41] was performed to evaluate antigen–antibody binding.

### 4.2 Statistical characterization of generative metrics

Critically, across five sequence-level metrics, inter-model variation was negligible (Table 3): diversity 0.507–0.516

(CV=0.8%), uniqueness 0.991–0.999 (CV=0.3%), novelty 0.925–0.977 (CV=2.2%), and Jaccard similarity 0.158–0.164 (CV=1.7%). The sole exception was perplexity, which showed substantial variation (range=14.14, CV=80.0%), with DeepSeek-V3-Ab achieving the lowest value (1.545) and Mistral-Ab the highest (15.681). Notably, DeepSeek-V3-Ab’s lowest perplexity coincided with the second-lowest novelty (0.930), suggesting a perplexity–novelty tradeoff warranting multi-seed investigation. As four of five metrics exhibited CV below 3%, architectural choice had minimal impact on generative quality at this scale, implying that generative capacity is primarily determined by model scale and training data rather than family-specific design elements. A Kruskal– Walli’s test found no significant difference in pLDDT distributions across the five transformer variants (Table 4) (H = 2.056, df = 4, critical χ^2^ = 9.488, p > 0.05). Pairwise Mann– Whitney U tests also showed no significant differences (all p > 0.18; smallest p = 0.185 for Llama-Ab vs. DeepSeek-Ab, Δmean = 0.87), indicating that at the 6-layer scale, transformer variant choice does not significantly affect structural quality of generated VH sdAbs. This result is consistent with the sequence-level metric analysis in Section 4.1 and reinforces the conclusion that, at the 6-layer scale employed here, the choice of transformer variant does not materially affect the structural quality of generated VH sdAbs. With N = 20 per group and SD ≈ 1.8 pLDDT, the test has ~80% power to detect mean differences ≥ 2.5 pLDDT. Observed differences were all < 1.1 (largest Δmean = 1.03, Mistral-Ab vs. Llama-Ab), indicating no practically meaningful differences between models, confirming all five compact variants generate stable, biologically plausible VH sdAb structures with means of 92.82–93.85 and 90–100% of sequences exceeding pLDDT ≥ 90, suitable for downstream analysis.

**Table 4:** Performance comparison of custom pre-trained compact transformer variants for novel VH sdAb generation.

| Model | Perplexity | Diversity | Uniqueness | Novelty | Jaccard Similarity |
| --- | --- | --- | --- | --- | --- |
| DeepSeek-V3-sdAbs | **1.545** | 0.509 | 0.997 | 0.930 | 0.164 |
| Llama4--sdAbs | 4.714 | 0.507 | 0.999 | 0.977 | 0.164 |
| Nemotron-sdAbs | 5.050 | 0.516 | 0.997 | 0.925 | **0.158** |
| Gemma3-sdAbs | 6.363 | 0.513 | 0.991 | 0.935 | 0.160 |
| AbLang-sdAbs | 7.001 | **0.783** | **1.000** | **1.000** | 0.060 |
| IgLM-sdAbs | 8.486 | 0.730 | **1.000** | **1.000** | 0.054 |
| Mistral-sdAbs | 15.681 | 0.507 | 0.999 | 0.945 | 0.164 |

Note: 1,000 generated samples per model based on OAS 15M HV samples training. Bold indicates best value per metric. **Perplexity** is a critical indicator of sequence naturalness in antibody design, where lower values reflect more biologically plausible and structurally coherent antibody sequences. AbLang, IgLM considered as baseline comparison.

**Table 4** shows that custom scratch fine-tuned models (Llama4-Ab, DeepSeek-V3-Ab, Mistral-Ab, Gemma3-Ab, Nemotron-Ab) generate more natural and biologically coherent VH sdAb sequences, reflected by low perplexity and strong control over diversity, uniqueness, and novelty. Among them, DeepSeek-V3-Ab achieves the best fluency (lowest perplexity = 1.545), followed by Llama4-Ab (4.714), indicating highly stable antibody-like sequence generation. In contrast, IgLM and AbLang show higher diversity and novelty but moderate quality, while NanoAbLlama performs poorly with extremely high perplexity (92.365), indicating unstable and less natural generation. Additionally, the custom compact fine-tuned models demonstrate strong structural reliability in terms of pLDDT, indicating high-confidence folding and reduced risk of misfolding, which is essential for accurate antibody design. Furthermore, novelty evaluation against a 15M training corpus confirms nearly 100% novel generation, with CDRH3 regions showing over 81% novelty compared to 15M, indicating that the models effectively capture biologically and chemically meaningful sequence patterns.

### 4.3 Case Studies (Ebola, HIV, HER2, SARS-CoV-2)

Pre-trained models were fine-tuned using disease-specific datasets, and training procedures were tracked using the Weights and Biases platform, as depicted in Appendix B - (b), (c), (d), and (e) for all disease targets. Fine-tuning adjusts the generative distribution toward disease-associated repertoire statistics and does not condition on antigen structure; generated candidates therefore resemble disease-associated antibodies statistically but are not guaranteed to bind the intended antigen

#### Ebola virus

For Ebola virus VH sdAb design, models were trained on 2,868 antibodies for 50 epochs and achieved 100% uniqueness and novelty. Diversity ranged 0.499–0.524 (mean 0.511 ± 0.009, CV 1.8%) and Jaccard similarity 0.132–0.153, indicating low overlap with known sequences. Perplexity varied, with DeepSeek lowest (2.093), suggesting high confidence. Overall, all compact transformer variants VH sdAb generation conditioned on Ebola-associated sequence repertoires related to Ebola (Appendix B, C).

#### HIV

For the HIV VH sdAb design, all fine-tuned models’ candidates generated from HER2-associated sequences with perfect uniqueness and novelty (1.0), indicating fully distinct and previously unseen antibodies. However, as noted in Section 3.1, the small dataset size (n = 430) means that perfect novelty may simply reflect insufficient data for memorization rather than better generative diversity. Diversity scores ranged from 0.572 to 0.711 (mean 0.665 ± 0.054, CV = 8.2%), with Llama4-Ab showing the highest diversity. The higher diversity observed for HIV compared to other targets reflects the broader sequence space explored when fine-tuning data provides weaker distributional constraints (Table 6, Appendix E).

**Table 6:**
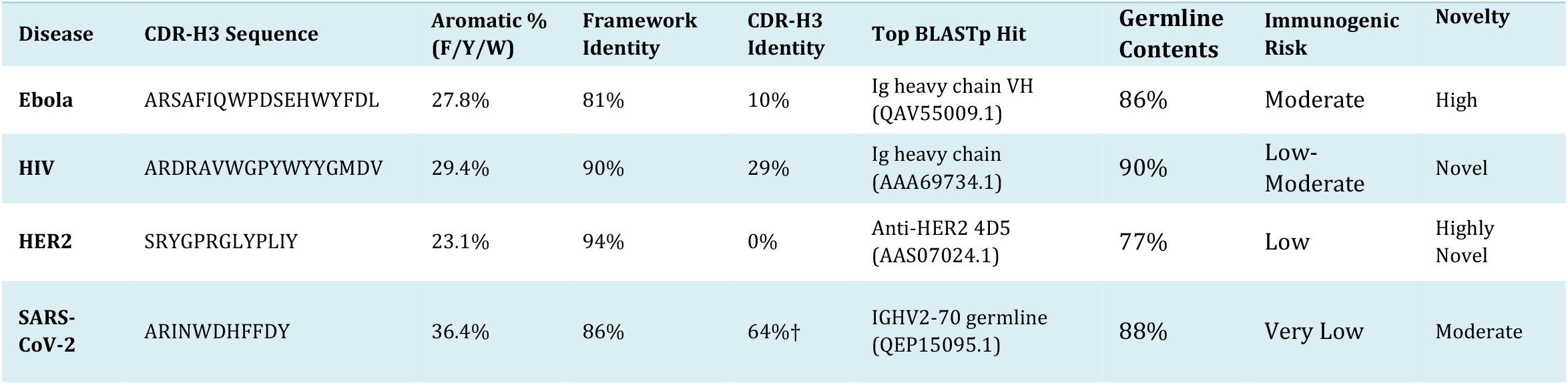
Biological Plausibility relevance, Immunogenic Risk analysis, and Sequence Novelty of LLM-Generated Antibody Candidates.

#### HER2

HER2 fine-tuning revealed a critical collapse to near-identical sequences (diversity ~0.065, Jaccard ~0.825) despite n = 22,778 training samples, likely due to the trastuzumab/pertuzumab-dominated low-diversity dataset. DeepSeek-V3-Ab avoided collapse (diversity 0.512, uniqueness 0.995, novelty 1.000, Jaccard 0.162), possibly due to MLA, though confirmation requires ablation and multi-seed studies. However, confirmation requires ablation and multi-seed studies, although other metrics remained outstanding.

#### SARS-CoV-2

For the COVID-19 VH sdAb design, Gemma3 exhibited the highest diversity (0.467) with perfect uniqueness and novelty (1.000) and low Jaccard similarity (0.187), suggesting effective generation of novel sequences, whereas other models produced slightly lower diversity but maintained high confidence in sequence generation (Appendix C). Diversity scores were comparable across models (mean 0.455 ± 0.008, CV = 1.8%).

### 4.4 3D Structure Evaluation on Fine-tuned models (AlphaFold-2, Boltz, RoseTTAFold-2, ESMFold)

Structural integrity was assessed by predicting 3D structures for the top 20 candidates per disease target using AlphaFold-2, with the highest-ranked sequence per target (by pLDDT) advanced to cross-platform validation via Boltz-2, ESMFold, and RoseTTAFold-2. All top-ranked candidates achieved pLDDT >80 and pTM >0.66, indicating well-folded, high-confidence structures. Cross-platform concordance was strong: mean pLDDT of 92.38±3.38 (Ebola), 90.76±3.52 (HER2), 91.16±3.97 (HIV), and 85.90±4.62 (SARS-CoV-2), all CV<6%. RoseTTAFold-2 RMSD values of 0.034–0.293 Å further confirmed structural consistency across independent folding algorithms (Table 5).

### 4.5 Docking, Developability, Humanness, and Immunogenicity Assessment

The selected novel VH sdAb sequences across all four disease targets exhibit desirable developability attributes and high structural stability (Appendix D). Sequence lengths ranged from 120 to 174 amino acids, with molecular weights between 13.2 and 19.3 kDa and moderately negative GRAVY scores (−0.264 to −0.311), indicating overall hydrophilicity.

#### Docking results

Docking was performed using HDOCK and HawDOCK against reference antigen structures (PDB IDs: 5JQ3 for Ebola, 3NGB for HIV, 1N8Z for HER2, 6MOJ for SARS-CoV-2; full results in Figure 3b and Appendix B). The Ebola candidate (Seq_2) showed the strongest predicted binding free energy (ΔG = −65.60 kcal/mol) and was computationally compared against the reference complex 5JQ3. When the natural heavy chain from the reference structure 3CSY was subjected to the same HDOCK/HawDOCK pipeline, it yielded a predicted ΔG of −49.56 kcal/mol, compared to −65.60 kcal/mol for our generated Seq_2 candidate (Appendix C, E). Notably, ΔG values are rigid-body docking estimates used for candidate ranking only and do not constitute experimental binding affinity measurements.

#### Humanness and immunogenicity analysis

To assess the biological plausibility and developability of the generated VH sdAbs, we performed a humanness evaluation [42] using ANARCI and IMGT numbering, germline V-gene assignment, and MHC-II immunogenicity profiling with NetMHCIIpan 4.3 against DRB1*0101 (Table 6). All four candidates passed the humanness evaluation with OASis identity of 92% (88th percentile) for the Ebola sequence and germline content ranging from 77–90%. CDR-H3 lengths of 13 amino acids are within the normal human VH range. Immunogenic risk, assessed by the number of strong MHC-II binding peptides, ranged from Very Low (SARS-CoV-2, 0 strong binders) to Moderate (Ebola, 4 strong binders).

#### VH sdAb design space and autonomous domain stability

The generated VH sequences are based on standard human germline genes (IGHV3-30, IGHV3-66, IGHV2-70) and may lack camelid nanobody hallmarks—hydrophilic substitutions at Kabat positions 37 (Val→Phe/Tyr), 44 (Gly→Glu/Gln), 45 (Leu→Arg/Cys), and 47 (Trp→Gly/Ser)—that enable stable, standalone VH domains. Their CDR-H3 lengths (11-17 aa) are shorter than typical nanobody loops (15–20+ aa), and the sequences likely retain hydrophobic VH-VL interface residues, risking aggregation or misfolding despite favorable in silico pLDDT scores. Future work should analyze key positions, incorporating nanobody-specific training or constraints.

**Figure 2:**
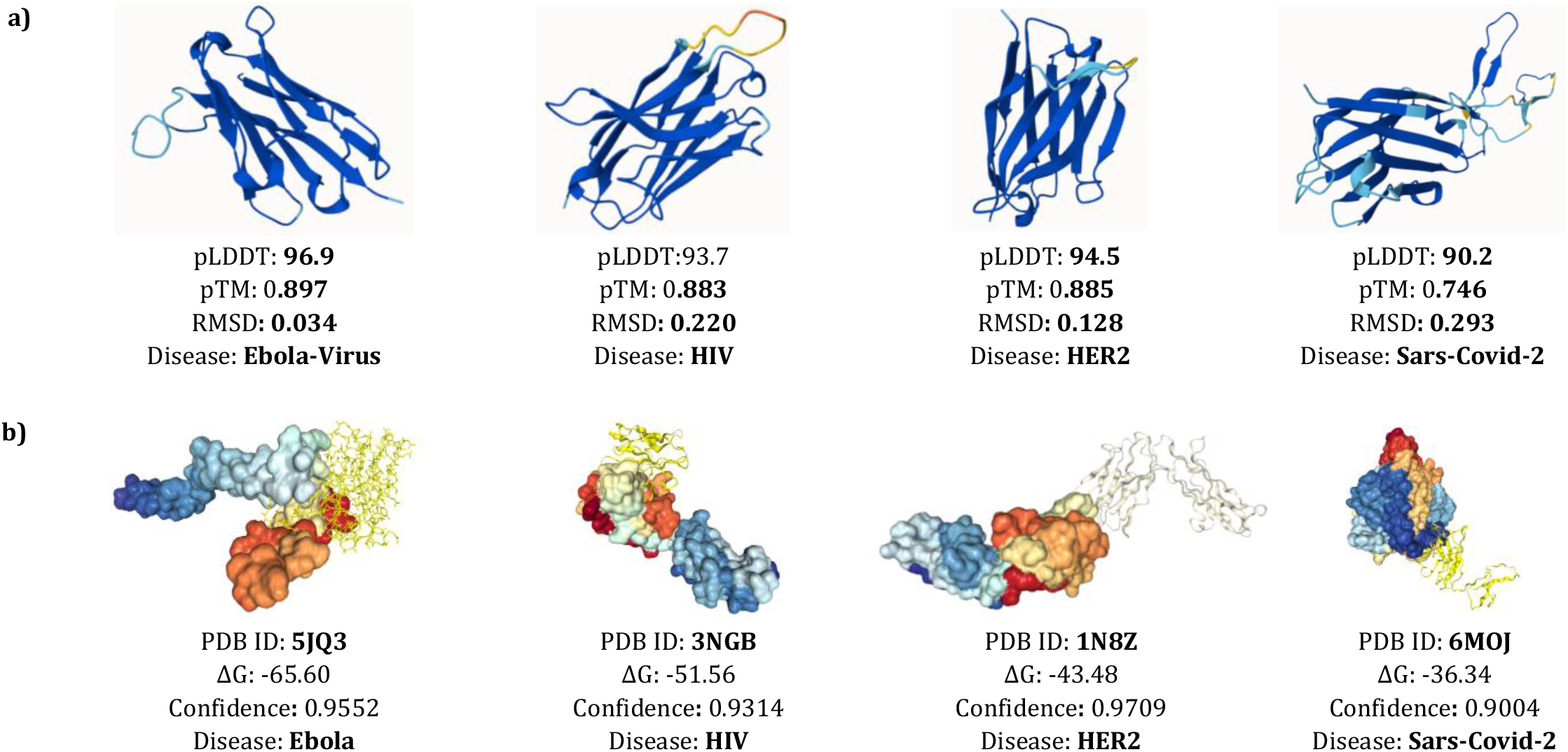
(a) Top-scoring 3D antibody structures selected from the best 20 sequences out of 1,000 generated candidates. (b) Sequences with the highest pLDDT scores were docked with their corresponding reference antigens (PDB IDs); the Ebola case shows the strongest binding free energy (ΔG ≈ −65).

## 5 Discussion

### Central Findings

All five compact transformer variants generated highly unique and novel VH sdAbs across four targets. They showed statistically indistinguishable structural quality (Kruskal–Wallis H = 2.06, p > 0.05; 80% power to detect Δ ≥ 2.5 pLDDT) and near-identical sequence metrics (diversity CV 0.8%, uniqueness CV 0.3%), indicating that generative capacity is driven by model scale and training data rather than architectural differences at this size.

### CDR-H3 Biological Plausibility Validation (ANARCI/IMGT, BLASTp Novelty)

CDR-H3 regions identified using ANARCI/IMGT and manually verified between the YYC motif and J-gene WGxGT anchor. BLASTp against NCBI nr showed framework identities of 81–94%, confirming human VH scaffolds. Germline content ranged from 77% (HER2) to 90% (HIV). CDR-H3 novelty was highest for HER2 (0% identity) and Ebola (10%), followed by HIV (29%), while SARS-CoV-2 had moderate identity (64%) due to its short 11-aa loop and rare IGHV2-7015 germline. Aromatic residues in CDR-H3 ranged 23.1–36.4%, exceeding natural averages (~20–25%), supporting antigen contacts. MHC-II profiling classified SARS-CoV-2 and HER2 as very low/low immunogenicity, with no strong binders in CDRs, confirming immunogenically silent binding surfaces (Table 6, Appendix F).

## 5 Limitations

All experiments used a single training seed (seed 42); multi-seed experiments (3–5 runs) are required to confirm whether the observed equivalence reflects true architectural insensitivity or single-seed stochastic convergence, as the statistical analyses quantify sequence variation rather than training variance. Structural evaluation covered only the top 2% of candidates (20/1,000), with one candidate per disease advanced to docking, representing a best-case pipeline. Fine-tuning adjusts distributional resemblance to disease-associated repertoires but does not condition on antigen structure, leaving binding selectivity computationally inferred. In addition, the diversity of disease-based studies under fine-tuning was low to moderate across all four case studies.

## 7 Conclusion

This study assesses new approaches by developing five compact generative transformer variants for de novo computational prioritization of VH single-domain antibodies across four disease targets (SARS-CoV-2, HIV, HER2, Ebola). The central finding is that at a compact scale (6 layers, ~2–10M parameters), architectural family choice has no statistically significant effect on generative quality (Kruskal–Wallis H = 2.06, p > 0.05), while training corpus diversity is the dominant factor, as starkly demonstrated by the HER2 mode-collapse failure. The compact variants produce VH sdAb candidates with high predicted structural confidence across multiple platforms (pLDDT consistently >80 across AlphaFold-2, Boltz-2, ESMFold, and RoseTTAFold-2) and favorable predicted biophysical properties, rigorous biological validation. However, these results characterize only the top 2% of candidates generated due to the time-intensive 3D structure prediction process. Additionally, the agentic evaluation pipeline demonstrates the feasibility of MCP-based integration with structure prediction platforms for automated antibody evaluation, establishing a design pattern that requires formal validation to assess its accuracy, throughput, and reliability.

## 8 ACKNOWLEDGMENTS

## Data Availability

The implemented code, all model weights, data, and agentic pipeline script will be open sourced upon review.

## Appendix

**Appendix A:**
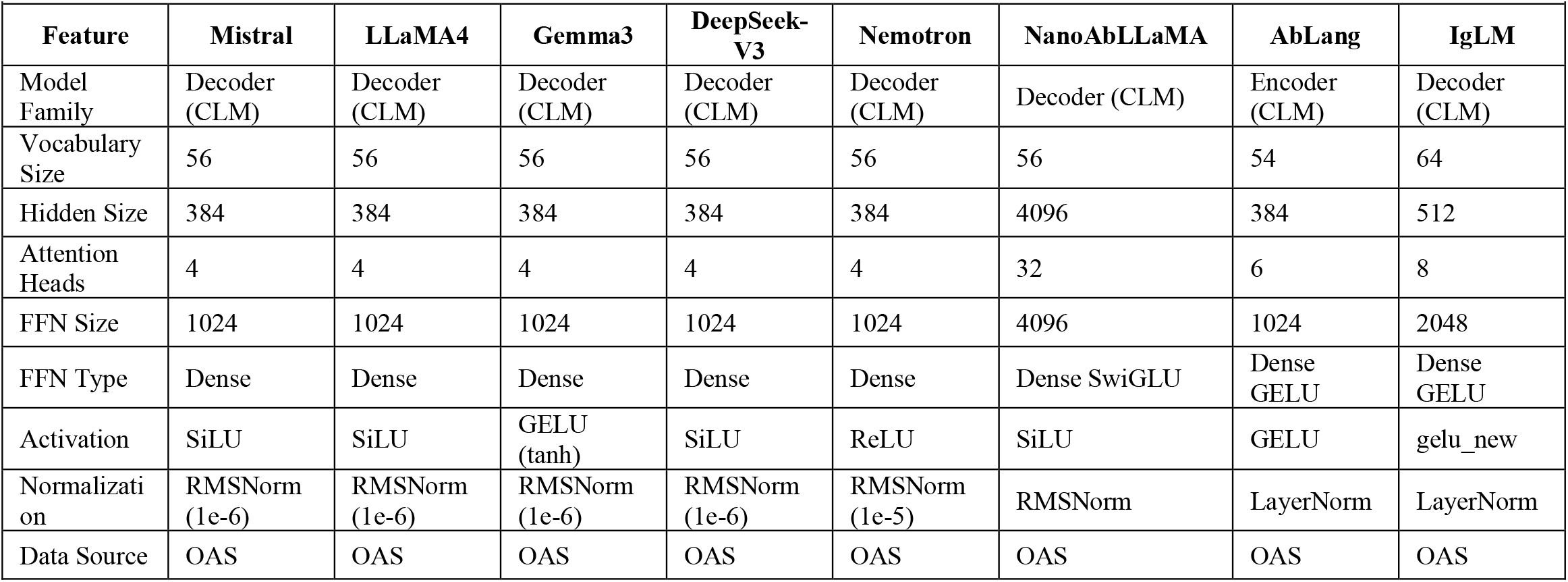
Training and Hyperparameters Configurations of Benchmark LLMs.

**Appendix B:**
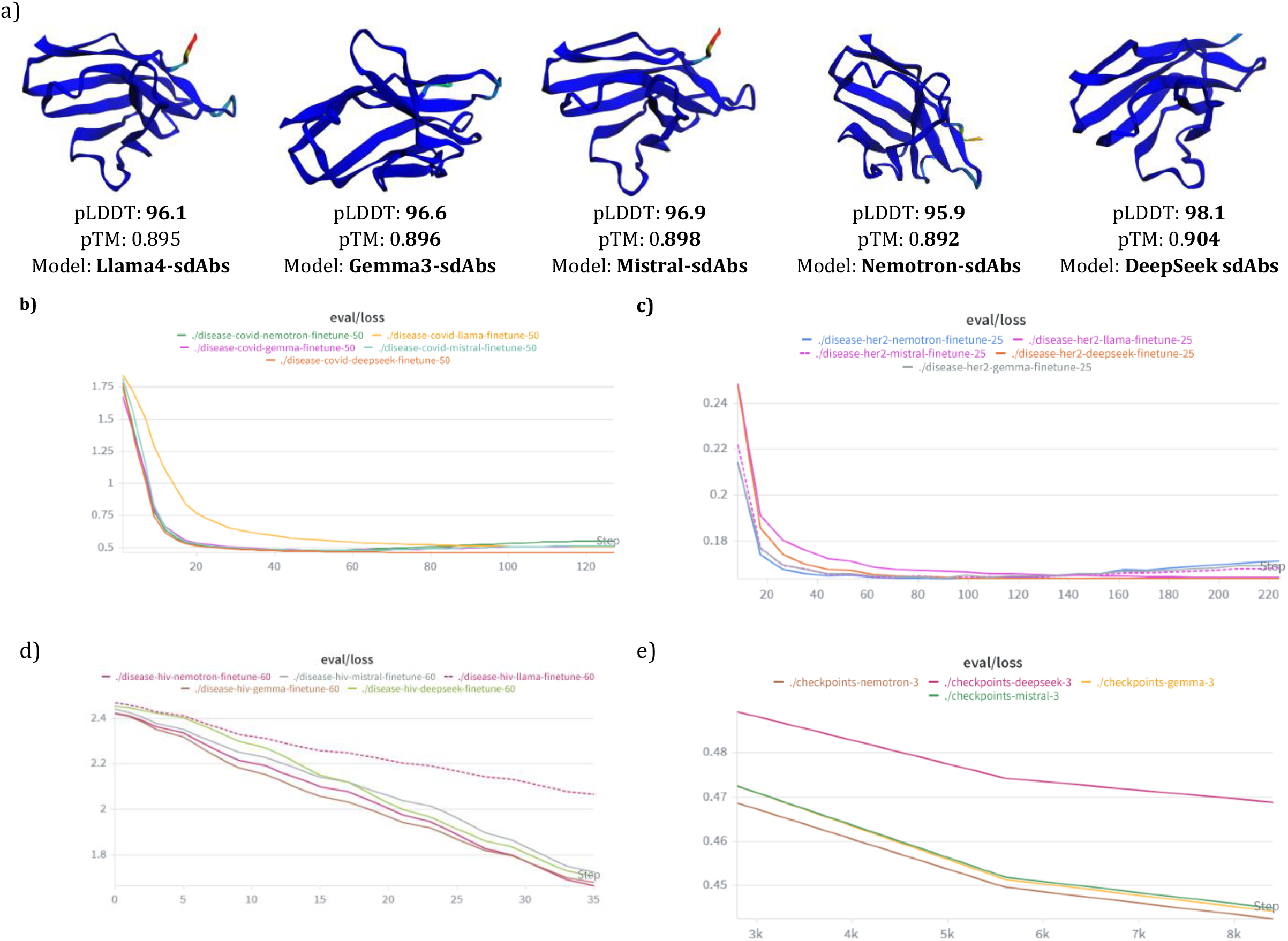
Illustration of 3D structures from pre-trained models (OAS Data) in **a)** and validation loss and training steps visualizing b, c, d, e are against Covid, HER2, HIV, and Ebola.

**Appendix C:**
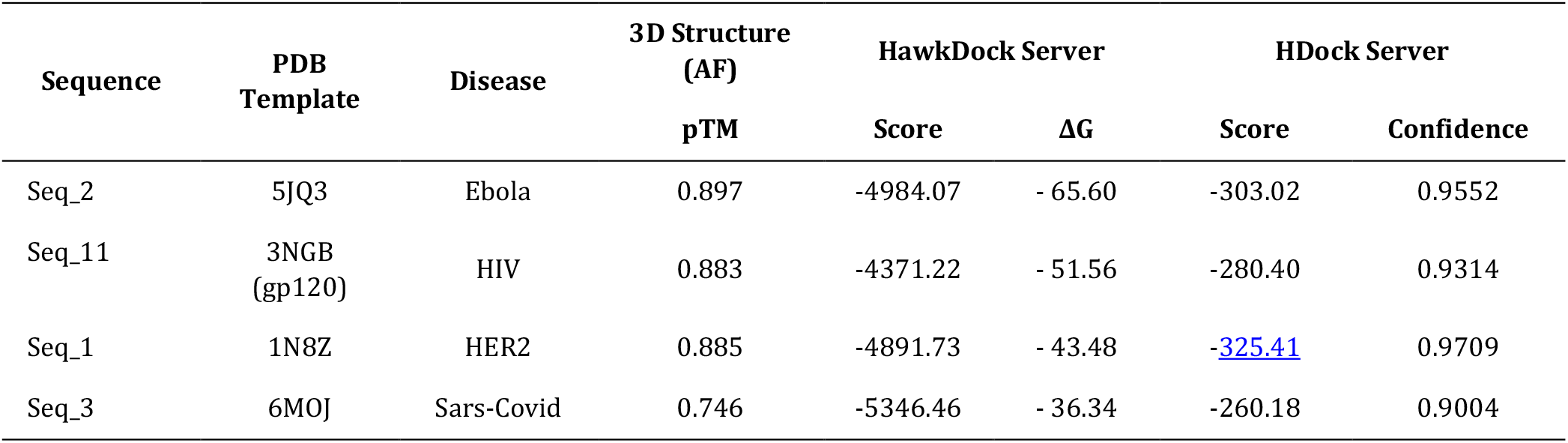
Antibody–Antigen Docking Results and best Structural validation Scores of four case studies.

| Sequence | PDB Template | Disease | 3D Structure (AF) | HawkDock Server |  | HDock Server |  |
| --- | --- | --- | --- | --- | --- | --- | --- |
| | | | pTM | Score | $\Delta G$ | Score | Confidence |
| Seq_2 | 5JQ3 | Ebola | 0.897 | -4984.07 | - 65.60 | -303.02 | 0.9552 |
| Seq_11 | 3NGB (gp120) | HIV | 0.883 | -4371.22 | - 51.56 | -280.40 | 0.9314 |
| Seq_1 | 1N8Z | HER2 | 0.885 | -4891.73 | - 43.48 | -325.41 | 0.9709 |
| Seq_3 | 6MOJ | Sars-Covid | 0.746 | -5346.46 | - 36.34 | -260.18 | 0.9004 |

**Appendix D:** Physicochemical (Developability) properties, & Docking results for top selected sequences.

| Seq_ID | Disease | Length | Weights | Gravy | deamidation_NS | Aromaticity | Agg. Hot | pH7 | $\Delta G$ |
| --- | --- | --- | --- | --- | --- | --- | --- | --- | --- |
| Seq_2 | Ebola | 125 | 14013.52 | -0.298 | 2 | 0.136 | 0.714 | 1.83 | - 65.60 |
| Seq_11 | HIV | 124 | 13795.40 | -0.311 | 2 | 0.145 | 0.857 | 5.79 | - 51.56 |
| Seq_1 | HER2 | 120 | 13194.71 | -0.264 | 1 | 0.125 | 0.857 | 3.79 | - 43.48 |
| Seq_3 | Sars-Covid | 174 | 19322.58 | -0.293 | 0 | 0.092 | 0.875 | 0.68 | - 36.34 |

**Appendix E:** HIV case study performance comparison of fine-tuned models using Transfer Learning.

| Model | Diversity | Uniqueness | Novelty | Jaccard Similarity | Perplexity |
| --- | --- | --- | --- | --- | --- |
| Llama4-HIV | 0.711 | 1.000 | 1.000 | 0.065 | 10.054 |
| Mistral-HIV | 0.694 | 1.000 | 1.000 | 0.077 | 12.477 |
| Gemma3-HIV | 0.676 | 1.000 | 1.000 | 0.089 | 10.114 |
| Nemotron-HIV | 0.674 | 1.000 | 1.000 | 0.083 | 6.373 |
| DeepSeek-V3 | 0.572 | 1.000 | 1.000 | 0.126 | 2.399 |

The Illustrated results are ordered based on the diversity of 1000 generated novel sequences.

**Appendix F:**
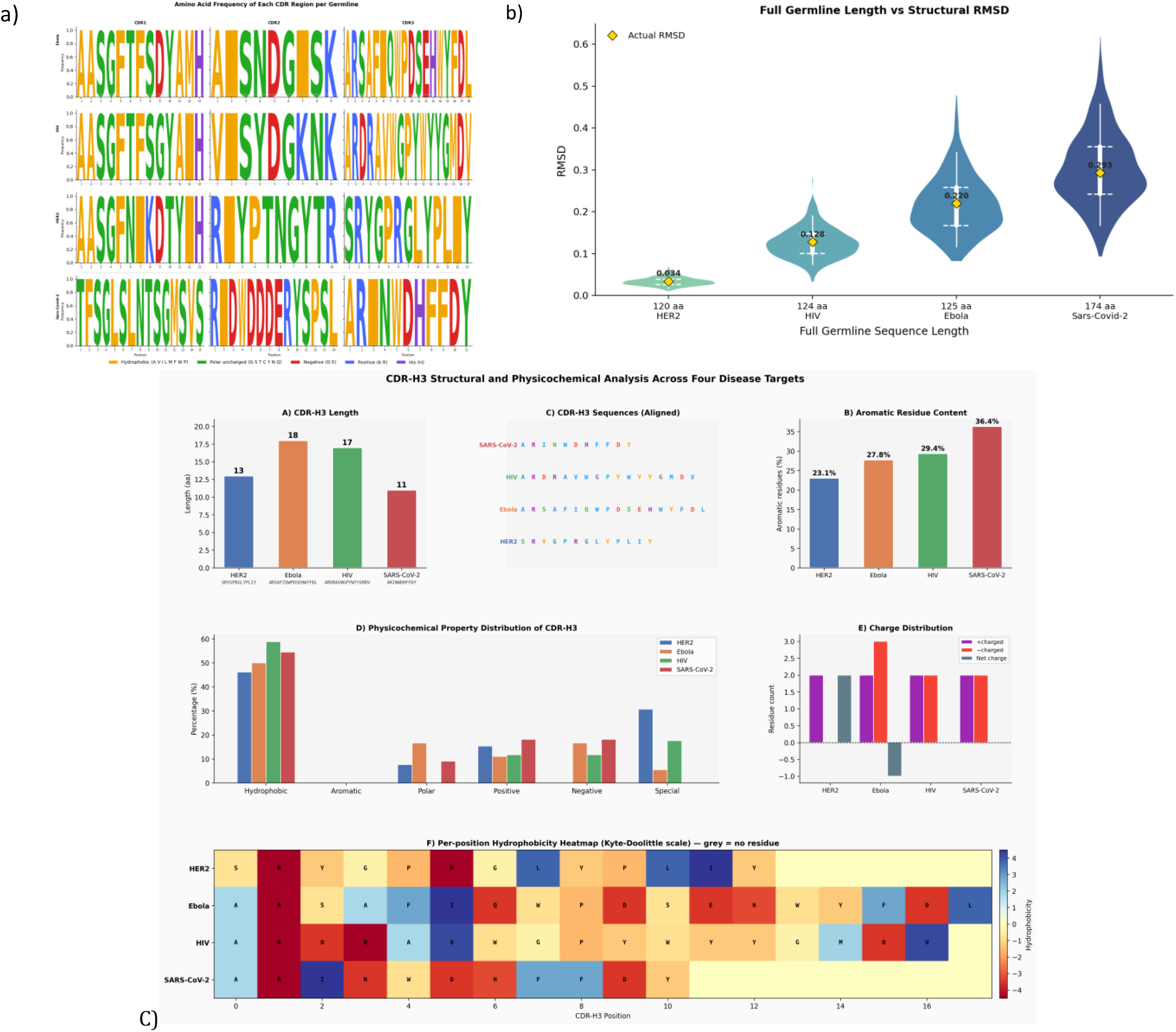
(a) Amino acid germline frequency and, repetition analysis across the four selected sequences. (b) RMSD comparison using RoseTTAFold-2, where the HER2 case shows the lowest structural deviation, illustrated in the violin plot, and C) CDRH3 distributio

**Appendix G:**
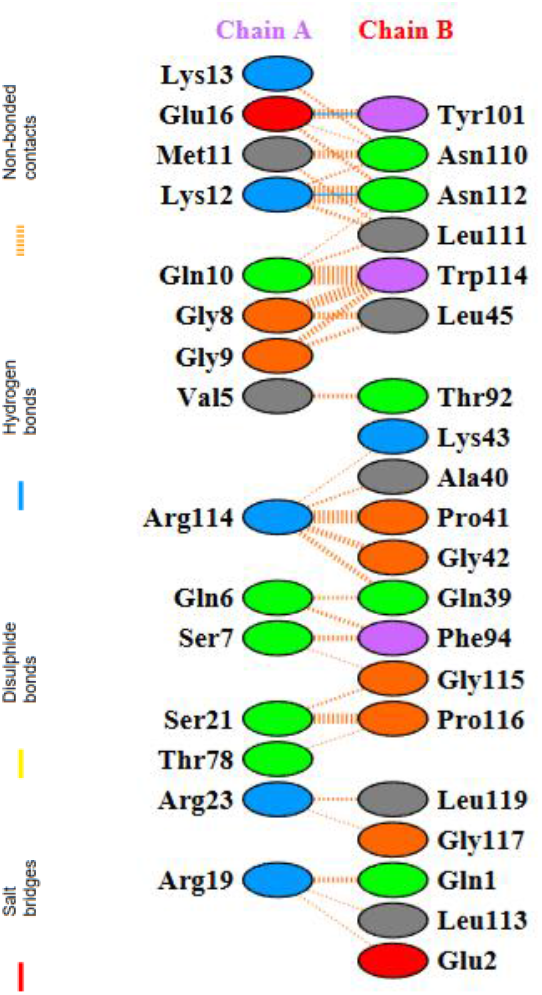
_Receptor (Antigen) and Ligand (Antibody) Residue Interaction of HIV (sequence id_11)

### Appendix H Formulation of Evaluation Metrics for Generated Antibody Sequences

#### 1. Diversity

Average pairwise dissimilarity between generated sequences:

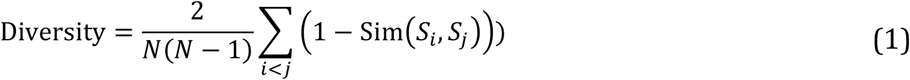

where (N) is the number of generated sequences and (Sim(*S*_*i*,_ *S*_j_)) is sequence similarity (e.g., normalized alignment identity or k-mer similarity).

#### 2. Uniqueness

Proportion of non-duplicate sequences among generated samples:

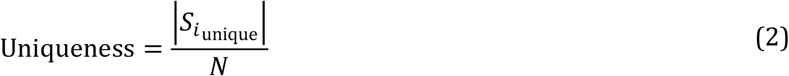

where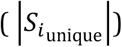 is the number of distinct sequences.

#### 3. Novelty

A fraction of generated sequences are not present in the training dataset (𝒟_*train*_):

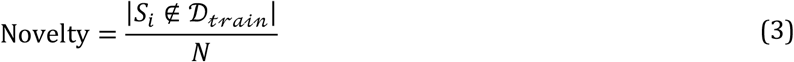

#### 4. Jaccard Similarity

Similarity between k-mer sets of generated sequence (S_g) and reference sequence (S_r):

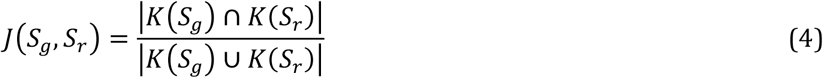

where (K(S)) denotes the set of k-mers extracted from sequence (S).

#### 5. Perplexity

Measure of language model uncertainty over sequence (S = (x_1, \dots, x_T)):

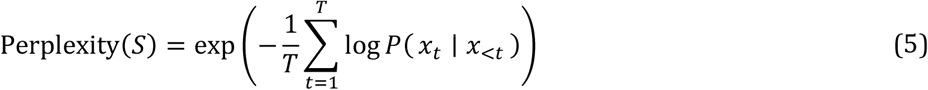

where (T) is sequence length and (*P*(*x*_*t*_ ∣ *x*_<*t*_)) is the predicted conditional probability.

### Appendix I Agentic assessment (Claude AI agent integrated Boltz-2, AlphaFold-2, ESM, RoseTTAFold-2)

- Confidence scores of the sequence length
- Residue and B-Factor analysis
- Sequence/ATOM Fragment profiling and analysis
- MCP, AlphaFold/RoseTTAFold/ESMFold connection with Claude Sonnet 4.6 Agent

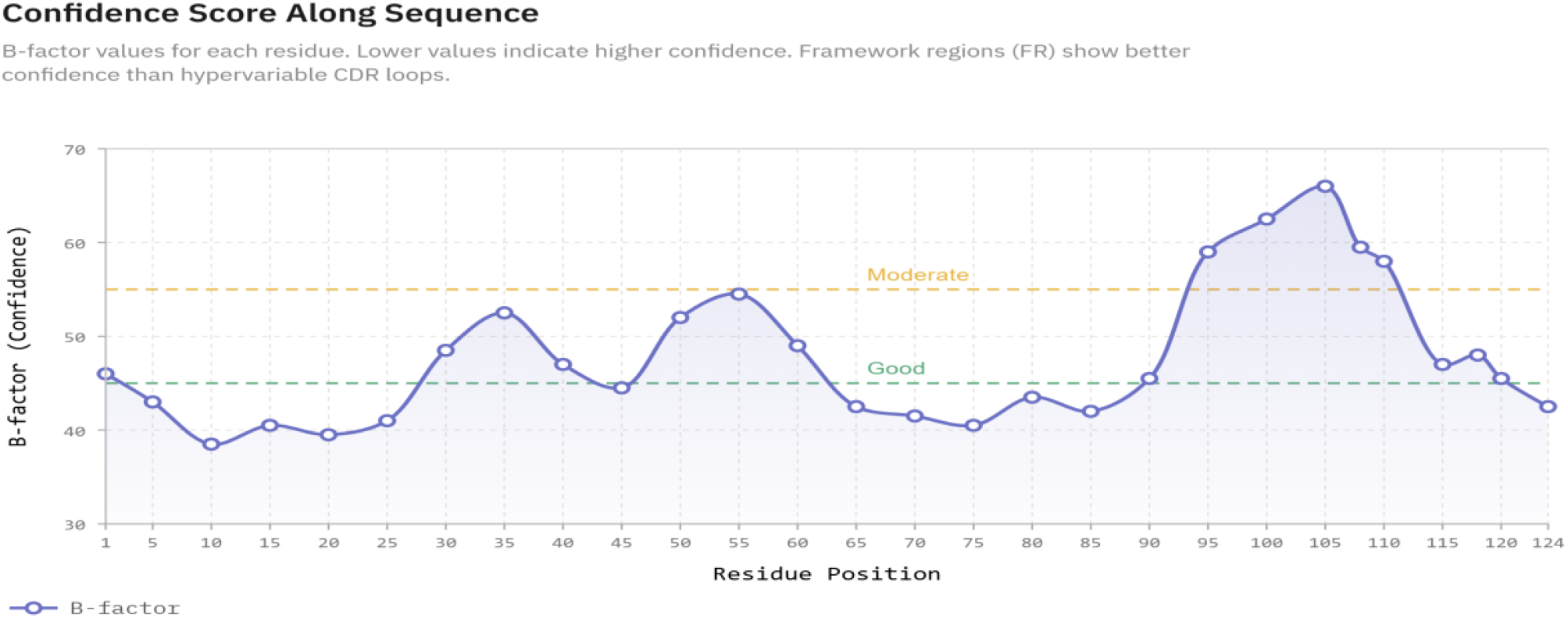

The table below illustrates residues and B-factor extracted from the Ebola sequence from the pdb complex by Claude AI, MCP integrated with Boltz-2 API

- Ebola Seq: QVRLVESGGGVVQPGRSLRLSCAASGFTFSDYAMHWVRQAPGKGLEWVTAISNDGISKYYADSVKGRFTISRDNSKNTLYLQMNSLRAEDT AVYYCARSAFIQWPDSEHWYFDLWGRGTLVTVSS
- Demo (Under Construction): https://claude.ai/public/artifacts/7683b38d-fe92-4377-9a88-40c547c27160

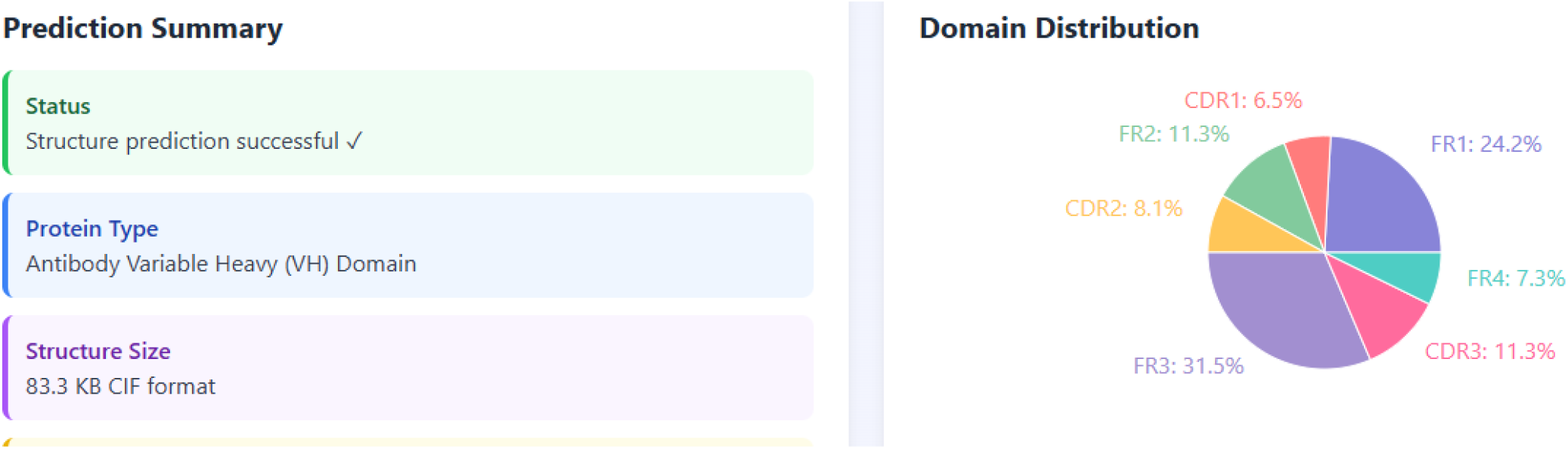

Illustrates each amino acid and its properties as sequence profiling for Ebola disease

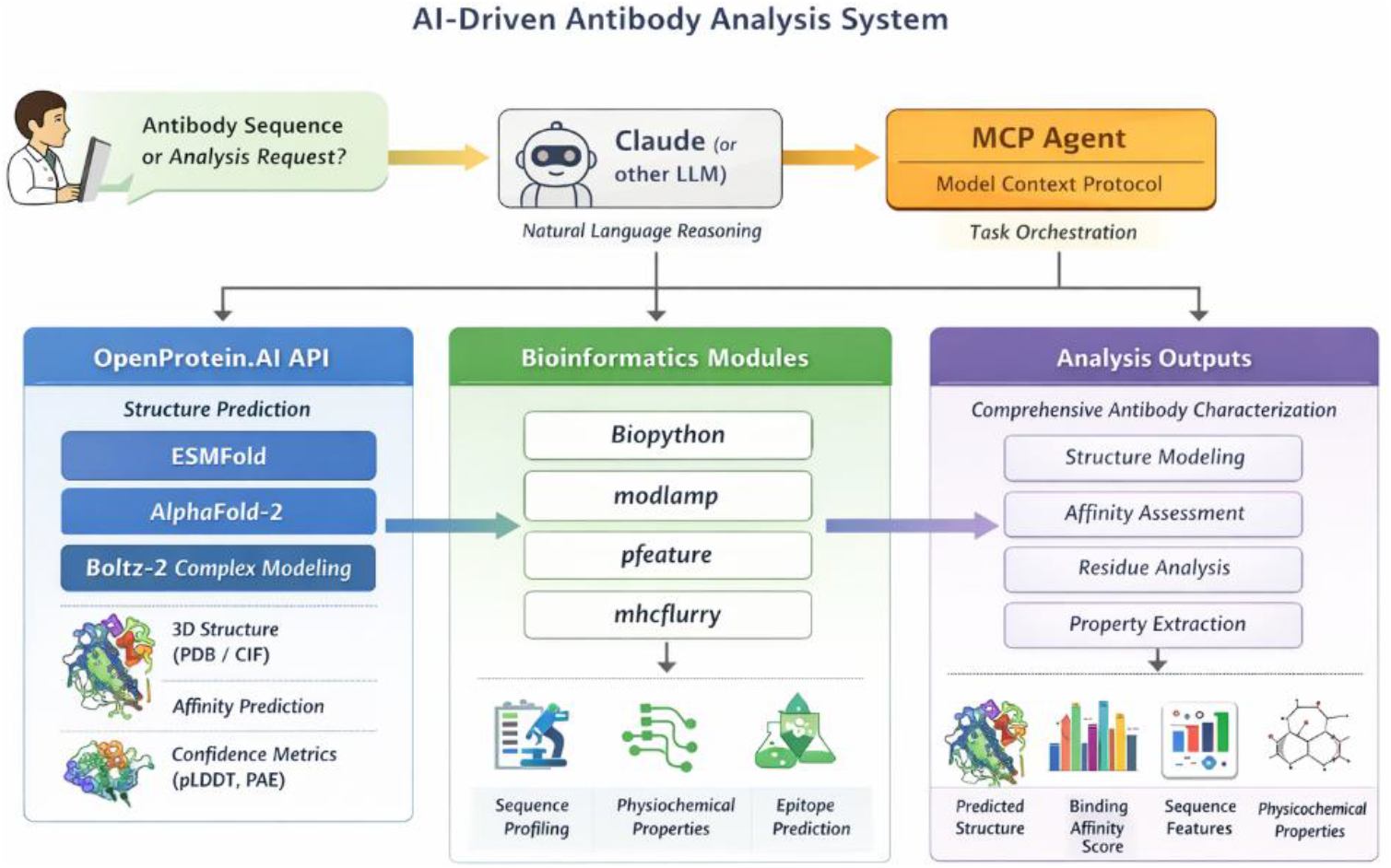

**Claude Sonnet 4.6, MCP (OpenProtein.AI) integrated with AlphaFold, Boltz-2, RoseTTAFold-2, ESMFold API**.

## Notes

### Competing Interest Statement

The authors have declared no competing interest.

